# IgE occupancy and antigen valency cooperate to control FcεRI aggregation geometry and signaling efficiency

**DOI:** 10.1101/2025.10.28.685118

**Authors:** Birgit Linhart, Rachel M. Grattan, Jon Christian David, Elton D. Jhamba, Christian Lupinek, Margarete Focke-Tejkl, Michael L. Paffett, Shayna R. Lucero, Rudolf Valenta, Lydia Tapia, Bruna Jacobson, Bridget S. Wilson, Diane S. Lidke

## Abstract

The crosslinking of IgE-bound FcεRI by multivalent allergens initiates mast cell and basophil signaling underlying Type 1 allergy. Yet, how allergen properties and IgE occupancy impact receptor aggregation and downstream signaling remain unclear. We used Phl p 1-specific IgE and recombinant fusion proteins presenting a Phl p 1-derived peptide in defined valencies and positions (MB1N, MB2N, MB4N, MB1N1C) to probe antigen-dependent signaling. Tetravalent MB4N evoked stronger degranulation and Ca2+ response than bivalent antigens, MB2N and MB1N1C. MB4N was also capable of signaling at low IgE occupancy and in Lyn-deficient cells. Monte Carlo simulations predicted that MB4N forms larger, complex receptor aggregates, while MB2N and MB1N1C produce dimers and linear chains. Consistently, addition of MB4N showed larger aggregates by electron microscopy and slower mobility by single particle tracking, compared to bivalent antigens. Thus, allergen valency and epitope spatial arrangement dictate FcεRI aggregate organization and subsequent effector cell activation.

## Introduction

Immunoglobulin E (IgE)-mediated allergies, including rhinitis, asthma, food allergy, atopic dermatitis, and life-threatening systemic anaphylaxis, affect a significant portion of the global population. These conditions diminish quality of life and cause a substantial burden on health care systems. Mast cells, which are central players in allergic responses, express the high-affinity IgE receptor, FcεRI, on their surface. Crosslinking of IgE-bound FcεRI by multivalent allergen triggers a cascade of signaling events that leads to mast cell activation and the release of inflammatory mediators. While the aggregation of IgE-FcεRI complexes represents the main pathomechanism of Type I allergies, much remains to be understood about the properties of allergens that generate an optimal effector cell response ^1^.

A number of structural parameters may influence signal initiation from FcεRI aggregates, including receptor spacing and orientation, as well as aggregate stability^2^. Steric constraints of certain configurations of allergen-IgE-FcεRIFc complexes have been attributed to the asymmetrical structure of bent IgE bound to FcεRI^3^ as well as to the distribution of IgE epitopes on allergens^1^. It has been well established that at least two IgE epitopes are required on an allergen to induce effector cell degranulation, and that increases in valency trigger even stronger responses ^4–8^. Paar et al used rigid, bivalent DNA spacers of varying length to show that spacing between receptors within an aggregate is a critical feature, with spacing as narrow as 40 Å capable of inducing degranulation^9^. Using a structurally defined trimer, Mahajan et al showed that the organization of the antigen-induced FcεRI aggregate can modulate the recruitment of positive and negative signaling molecules, which in turn controls the type of signaling propagated^10^. In addition to spacing and valency, the affinity of IgE for the antigen has also been shown to modulate FcεRI signaling outcomes ^11–14^. Ligands with low affinities to IgE were shown to form complexes with shorter lifetimes that would result in more transient aggregation^11^. Studies comparing low- and high-affinity ligands indicated that signaling differences were due at least in part to differential recruitment of membrane scaffolding proteins, LAT versus NTAL^14^. Finally, it is also important to consider the number of antigen-specific receptors available for crosslinking. It is estimated that less than 10% of the IgE found on a single cell are specific to the same antigen^15^. Therefore, the size of aggregates formed is restricted by the repertoire of IgE idiotypes bound to FcεRI on the mast cell surface.

A limitation of most studies is the lack of defined molecules reflecting the natural IgE response, which is mainly directed to protein antigens in Type I allergies^16^. In early signaling studies, bivalent crosslinking using either anti-IgE or anti-FcεRI antibodies was a common approach ^17,18^. Antigen mimics are also useful, often based upon random conjugation of small synthetic haptens and the corresponding hapten-specific IgE; the widely used model system of DNP/anti-DNP IgE is an important example of this strategy^19^, but is limited by lack of structural information with regards to epitope valency and spacing. We recently developed a molecular model system which is based on the engraftment of an IgE epitope derived from the major timothy grass pollen allergen, Phl p 1, onto a monomeric scaffold protein, myoglobin^20^. Several allergenic molecules were produced as recombinant proteins, each with a defined number and position of IgE epitopes. Using a monoclonal Phl p 1-specific IgE, it could be shown that number and position of IgE epitopes on the recombinant protein affected 1) the formation of allergen-IgE immune complexes in solution, 2) the *in vitro* degranulation of effector cells and 3) allergic inflammation *in vivo*. It was demonstrated that as many as four IgE antibodies could bind in close proximity to an epitope, inducing strong mast cell activation. In the present study, we used these structurally defined antigens to test the hypothesis that the number of epitopes present on an allergenic molecule and their spatial arrangement influences the degree of FcεRI aggregation and the subsequent signaling cascade. We focus on understanding the link between the geometry of receptor aggregates and signaling outcomes, both as defined by antigen epitope spacing and valency, as well as the number of antigen-specific, IgE-bound FcεRI available for crosslinking. Receptor aggregation was assessed by high resolution imaging techniques, including single particle tracking and transmission electron microscopy. Results were integrated into a mathematical model for a better understanding of the impact of receptor aggregation on signaling.

## Materials and Methods

### Monoclonal IgE antibody, recombinant proteins

A hybridoma cell line secreting a mouse monoclonal IgE antibody specific for the peptide aa 86- 116 (EPVVVHITDDNEEPIAPYHFDLSGHAFGAMA) derived from the major timothy grass pollen allergen, Phl p 1 (UniProt P43213), was established by immunization of BALB/c mice with the KLH-coupled peptide ^21^. Hybridoma cells were cultivated in CD Hybridoma medium (Gibco, Thermo Fisher Scientific, Waltham, MA, USA) for antibody production. The monoclonal IgE antibody (mIgE) was purified from the hybridoma supernatant by affinity chromatography using a 1 mL HiTrap NHS-activated HP column (GE Healthcare, Chicago, IL, USA) loaded with 1 mg of the Phl p 1-derived synthetic peptide according to the manufacturer’s instructions, and dialyzed against PBS pH 7.4. Myoglobin derivatives containing 1-4 copies of the Phl p 1-derived IgE-reactive peptide DLSGHAFGAMA were produced as previously described ^20^. Briefly, synthetic genes (ATG Biosynthetics, Merzhausen, Germany) encoding MB1N, MB2N, MB1N1C, and MB4N along with a C-terminal 6x histidine tag were cloned in the expression plasmid pET17b (Novagen, Millipore Sigma, Burlington, MA, USA). Recombinant proteins were expressed in *E.coli* BL21 (DE3) cells (Agilent, Santa Clara, CA, USA) after induction with 1 mM IPTG and subsequent cell culturing for 4 h. Proteins were found in the water insoluble fraction of bacterial lysates and purified on a Ni-NTA column under denaturing conditions according to the manufacturer’s instructions (Qiagen, Hilden, Germany). Proteins were dialyzed against 10 mM NaH_2_PO_4_, pH 8.0, and purity of the proteins was checked by SDS-PAGE. Protein concentrations were determined by Micro- BCA protein assay (Pierce, Rockford, Ill, USA). Myoglobin from equine heart was purchased from Sigma-Aldrich (Vienna, Austria).

### Degranulation assay

Rat basophilic leukemia (RBL-2H3) cells were grown in 96 well tissue culture plates for 24 h in MEM (Life Technologies, Grand Island, NY) supplemented with 10% fetal bovine serum, 1% penicillin-streptomycin and 1% L-glutamine (Invitrogen, Thermo Fisher Scientific, Waltham, MA, USA) as described ^22^. RBL-Lyn^KO^ cells, which were previously generated by CRISPR-Cas9 gene editing, were cultivated under the same conditions as the WT cells ^23^. Cells were passively sensitized with Phl p 1-specific mIgE (2 µg/mL or as indicated) for 2 h, washed with Hanks’ balanced salt solution (HBSS), and stimulated in HBSS with the indicated concentrations of recombinant myoglobin derivatives or, for control purpose, myoglobin, or buffer for 30 min at 37°C. Degranulation of cells was quantified by measuring ß-hexosaminidase release in supernatants as previously described ^24^. Percent ß-hexosaminidase release was calculated by cell lysis with 1% Triton X-100, and fraction of 100% release are shown. All measurements were performed in triplicates.

### Quantification of mIgE binding to FcεRI on RBL-2H3 cells

A fluorescent IgE conjugate was created by labeling the Phl p-1 specific monoclonal IgE antibody with Alexa Fluor 647 (Invitrogen; Carlsbad, CA) according to the manufacturer’s instructions, whereby a dye:IgE molar ratio of either 0.96:1 or 3.63:1 was achieved. RBL-2H3 cells were incubated with the mIgE^Alexa-647^ antibody for 2 h in different concentrations (Supplementary table S1) and washed twice to remove unbound IgE. Cell-associated fluorescence intensity was measured for each concentration on a FACS Calibur (BD Biosciences) and assigned in molecules of soluble fluorochrome (MESF) units using the Quantum Alexa Fluor 647 MESF calibration kit (Bangs Laboratories, Fishers, IN) according to the manufacturer’s instructions. Numbers of mIgE^Alexa-647^ bound to surface FcεRI at each concentration were calculated based on the established standard curve and the degree of labeling of the IgE and related to the ß-Hexosaminidase-release in RBL-2H3 cells primed with the same IgE concentrations and activated with 1 nM of myoglobin derivatives or only myoglobin.

### Determination of kinetics and binding affinities by surface plasmon resonance measurement

Surface plasmon resonance measurement was performed as a single cycle kinetic on a Biacore 2000 (GE Healthcare, Uppsala, Sweden) at 25°C using a CM5 sensor chip (GE Healthcare). The surface was activated with a 1:1 mixture of 1-ethyl-3-(3-dimethylaminopropyl carbodiimide) hydrochloride and N-hydroxysuccinimide at a flow rate of 5 μL/min for 7 minutes. An anti-mouse IgE antibody (BD OptEIA™ Mouse IgE ELISA Set, BD Biosciences, San Jose, CA, USA) was resolved in 10 mM sodium acetate pH 4.5 according to the pH scout and immobilized as a capturing antibody to a maximal response of 100 RU (resonance units). The chip surface was inactivated by injecting 1 M ethanolamine-HCl (pH 8.5) followed by saturation of the surface with MB1N and loading with monoclonal IgE at a flow rate of 5 µL/min for 3 min. The reference cell was not loaded with mIgE. The following concentrations of MB1N were injected for the single cycle kinetic: 0.333, 1, 3, 9, 27 nM with a flow rate of 30 µL/min. Dissociation of MB1N was investigated by injecting HBS-EP for 30 minutes at 30 μL/min. No dissociation of mIgE was observed 30 min after the highest MB1N concentration. Dissociation constants and rate constants (on-rate, off-rate) were calculated with BIA evaluation software 3.2 (GE Healthcare) using a 1:1 interaction model.

### Fura-2AM calcium assay

Calcium release in mIgE-loaded RBL-2H3 cells after activation with MB2N, MB1N1C, or MB4N was measured in a Fura-2 AM calcium assay. For this purpose, RBL-2H3 cells were primed with 2 µg/mL mIgE in supplemented MEM for 2 h, washed with HBSS and incubated with 2 µM Fura- 2 AM (Molecular Probes, Eugene, OR) solution for 30 min at RT. Cells were washed with HBSS and rested for 20 min in 37°C incubator. As a negative control, unprimed cells were used. Ratiometric measurements were performed at 35°C using an Olympus IX71 inverted microscope outfitted with a UPLANSAPO 60X/NA1.2 water emersion objective coupled to an objective heater (Bioptechs). After 30 s, cells were activated with MB1N, MB2N, MB1N1C, or MB4N and ratiometric changes in cytosolic calcium were monitored for a total of 5 min. Alterations between 340 and 380 nm at 1 Hz were done using a xenon arc lamp monochromator (Cairn Research OptoScan). Fura-2 fluorescence emissions at 510 nm were measured with an iXon 887 electron-multiplying charge-coupled device (EMCCD) camera using IQ3 imaging software (Andor Technology). A previously described in-house MATLAB script was used for ratiometric analysis for each cell (15–20 cells per field of view) ^25^.

### Transmission Electron Microscopy

Apical membrane sheets were prepared using a “rip-flip” technique as previously described ^26^. RBL-2H3 cells were plated on glass coverslips at 70% confluency and primed overnight with 1 µg/mL Phl-P1 IgE. Where indicated, cells were treated with 1 nM of bivalent or tetravalent antigens in HBSS for 5 min at 37°C. The cells were briefly fixed and the glass coverslips were inverted onto formvar and carbon coated, poly-l-lysine treated nickel grids (Electron Microscopy Sciences). Pressure was applied to adhere the apical cell membranes to the grids, and the membranes were ripped from the cell as the coverslip was removed. Membrane sheet laden grids were fixed with 2% paraformaldehyde and labeled with anti-FcεR1β antibody (Santa Cruz Biotechnology) and 6 nm colloidal gold conjugated secondary antibody (Jackson ImmunoResearch). The samples were postfixed with 2% glutaraldehyde and stained with 0.03% tannic acid and 2% uranyl acetate. Digital images were acquired on a Hitachi 7700 Transmission Electron Microscope. Images were analyzed with Image J (NIH) and MATLAB (Mathworks).

### Single Particle Tracking

Single particle tracking (SPT) was performed using RBL-2H3 cells stably expressing an FcεRI γ-subunit tagged at the N-terminus with an HA-tag and a HL4.1 fluorogen activating peptide (FAP), previously characterized in Schwartz 2015 ^27^. HA-QD was prepared as described in Valley 2015 ^28^.

### Mathematical modeling

#### Antigen PDB model construction

We used artificial allergen constructs defined in ^20^ to construct all-atom models (PDB structures) for MB1N, MB2N, MB1N1C, and MB4N. The primary structures were input into RosettaFold ^29^ and AlphaFold2 ^30^. For each structure, six models were produced, one from AlphaFold2 and five from RosettaFold. We used UCSF Chimera ^31^ to add hydrogens and charges to these models before performing energy minimization using AMBER ff14SB. During minimization there were 5 iterations of 100 gradient descent steps, where the steepest gradient descent step is 0.02 Å. On average, at the end of energy minimization, the energy difference between steps is on average 204.48 kJ/mol, with standard deviation of 82.95 kJ/mol. The same IgE model from ^32^ was used.

#### Antigen PDB model selection

We selected a representative subset of the PDB structures produced from the model construction process. For MB1N and MB2N, we selected the constructs with the lowest energy values after minimization. For MB1N1C two energetically similar PDB structures with low energies -12613.35 kJ/mol and -12159.14 kJ/mol were found to have distinct conformations; the RMSD between the two models is 0.52 Å. These versions, MB1N1C (closed) and MB1N1C (open), differ in accessibility with respect to their C-termini. Using relative solvent accessibility surface area (relSASA) for MB1N1C (closed) and MB1N1C (open)’s C-terminus were calculated in Chimera, resulting in an average relSASA of 0.40 for MB1N1C (closed), with standard deviation σ=0.34, and 0.59 for MB1N1C (open) with standard deviation σ=0.26. These values suggest that MB1N1C (closed)’s C-terminus epitope may be less accessible. Finally, models obtained for MB4N were very different in RosettaFold and AlphaFold2, which may indicate that the long tail containing 4 epitopes is disordered. Therefore, the model chosen for MB4N was an energy minimized model where all four epitopes have relSASA values near 1.0; it has a post-minimization energy of –10804.46 kJ/mol.

#### Monte Carlo simulation of IgE-Antigen aggregation

We used Monte Carlo-based simulation to geometrically model the diffusion and aggregation of IgE and artificial allergens. At each timestep, individual molecules diffused on a 200 x 200 nm^2^ planar surface representing a patch of a cell surface with diffusion coefficient 0.09 μm^2^/s ^8^. Aggregates diffuse more slowly by a factor of (1/N), where N is the number of IgE in an aggregate ^32^. Simulation setup parameters and 3D iso-surface model generation is described in ^32^. Resampling was done whenever molecules collided, or when molecules were outside the bounds of the patch. Binding between an IgE and an allergen can only occur when both have at least one unbound binding site; those two binding sites will bind if the Euclidean distance between them is less than 17.34 Å, to allow for some modeling of flexibility in a strictly rigid-body simulation. At any given timestep, any pair of bound binding sites has an unbinding probability of 10^-5^. Experiments were run for a ratio of 1:1 IgE to antigen molecules. There were 14 IgE molecules in the simulation, consistent with a density of 350 IgE molecules/μm^2^ when the cell surface is primed with 110,000 IgE molecules. The simulation terminated when 500k timesteps finished or when resampling failed 1000 times, consecutively.

#### Rule-Based Modeling

We simulated the formation of receptor aggregates on a cell surface by performing a rule-based model simulation using the BioNetGen extension for Visual Studio Code v.0.7.2 9 ^33^. The model captured the formation of aggregates of any size and crosslinking can be observed. The total number of receptors in aggregates as a function of time was counted. We ran simulations for the following numbers of IgE molecules: 2155, 3795, and 14830. Number of ligands in solution was kept constant at 20,000 molecules. For a 1 nM solution, there were 1.38 x 10^6^ ligands per cell, which was far in excess of the number of available receptors, thus keeping the background number of ligands fixed even as more ligands bound to the cell surface was justifiable. Simulations use the experimentally found values for k_off_ = 1.72x10^-3^ s^-1^ and K_D_ = 8.95x10^-10^ M. The number of cells per well was 20,000. For both MB4N and MB2N, the k_on_ for all binding sites was identical. For MB1N1C, to emulate the partially hidden position of the epitope in the C terminus, we used the results from the geometric model to estimate the relative on-rate at the C terminus compared to the on-rate at the N-terminus epitope. In the geometric models we observed that in events where only one epitope was bound, the N-terminus is bound 83% of the time, while the C-terminus is bound 17% of the time. By assuming rates are directly proportional to binding probabilities, we can assume that the on-rate at the C-terminus epitope is nearly 5-fold lower than the on-rate at the N-terminus. For comparison, we found that in MB2N the two epitopes are bound with equal probability, so use of the same on-rate is a reasonable assumption. Crosslinking binding rates are calculated from Eq.(2) in ^34^: ß= K_x_C_R_ assuming the dimensionless constant ß =1, where K_x_ is the crosslinking equilibrium rate, C_R_ is the concentration of receptors, and that the crosslinking off-rate has the same value as the experimental off-rate. Calculations were performed using stochastic Network-Free simulations, and each time series was repeated 5 times. Error bars indicate the standard error of the mean. Simulations were run from t=0s to t=300s (5 min) in steps of 20 seconds.

## Results

### Availability of Phl p 1-derived epitopes in engineered antigens

A previously developed molecular model system based on a peptide derived from one of the most relevant respiratory allergens worldwide, the grass pollen allergen, Phl p 1, and a corresponding mouse monoclonal IgE antibody, was employed to study allergen-induced FcεRI aggregation and subsequent effector cell activation^20^. As shown in Figure 1A, a monomeric scaffold protein, myoglobin, was engrafted at the N-terminus with one, two, or four copies of the IgE-reactive peptide (colored blocks delineated with the letter P), resulting in the engineered molecules MB1N, MB2N, MB1N1C, and MB4N. The pair of molecules with a valency of two, MB2N and MB1N1C, differ in epitope placement; for MB2N, the IgE-binding sites are in close proximity while MB1N1C contains two peptides separately located at the N- and C-terminus. The four myoglobin derivatives were produced as recombinant proteins in *E. coli* and purified via a C-terminal 6x-histidine tag as previously described^20^. We determined the affinity of the monoclonal peptide-specific IgE antibody to the monovalent MB1N by surface plasmon resonance (SPR) measurements (Fig. 1B). The sensorgram captures mIgE binding to increasing concentrations of MB1N in a single cycle kinetic. Recorded (black) and calculated (red) values were superimposed using the 1:1 interaction model, showing an excellent fit of the 2 curves. We chose the 1:1 interaction model because, despite the bivalent nature of the monoclonal IgE, the MB1N antigen presents only a single epitope, allowing each binding event to be accurately described as a simple 1:1 interaction. We found that the mIgE antibody bound MB1N with high affinity (dissociation constant K_D_: 8.95x10^-10^M). While the association rate constant (k_a_) of 1.91x10^6^ M^-1^s^-1^ suggests an initial strong complex formation, the dissociation rate constant of 1.72x10^-3^/s corresponds to an estimated half-life of only 6.7-8 min. These values are used for parameterization of antigen binding in the mathematical models below.

**FIGURE 1.**
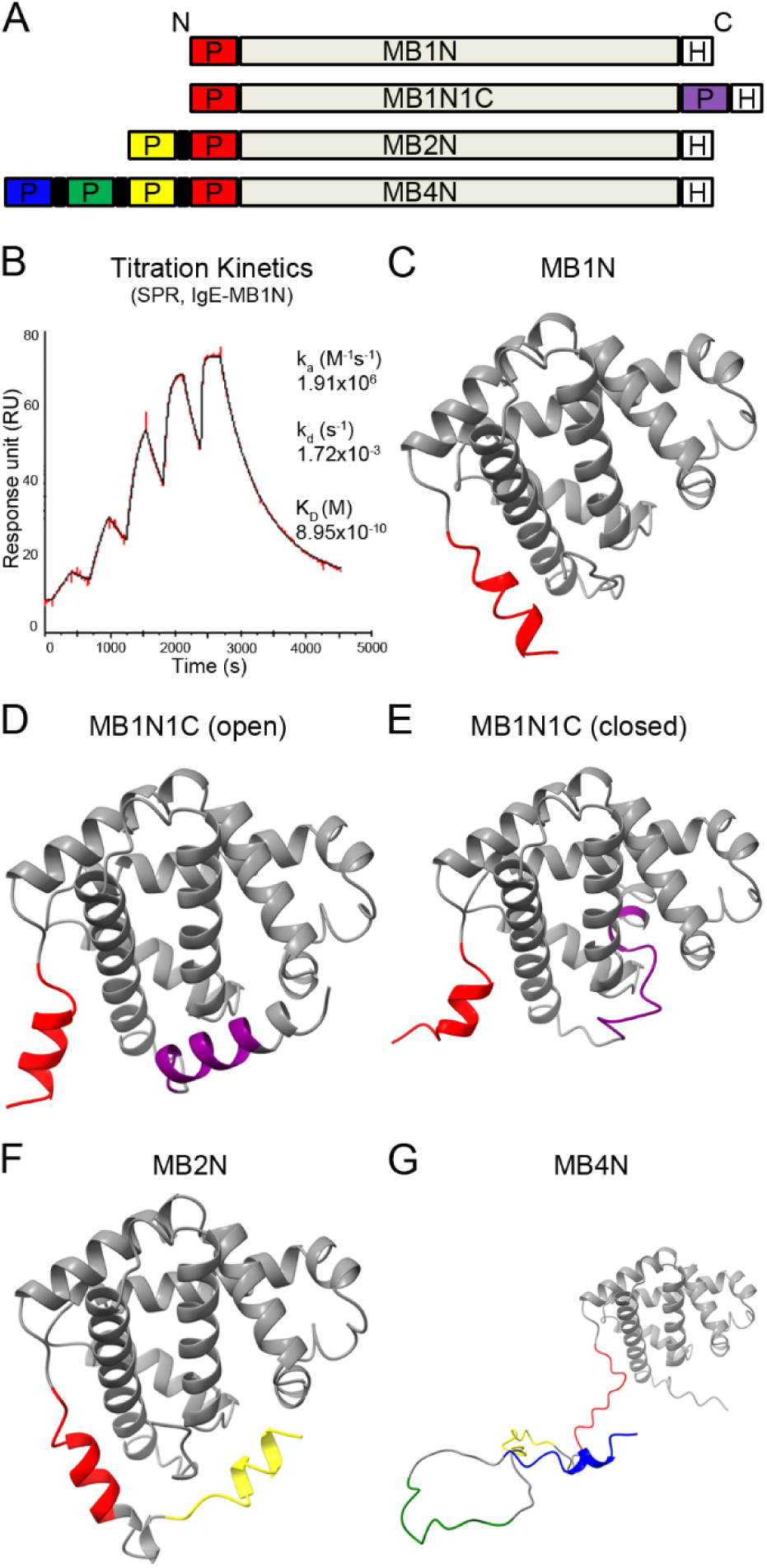
Structural design of engineered IgE-binding antigens. (*A*) Schematic representation of the myoglobin backbone (*grey*) engrafted with different numbers of a Phl p 1-derived IgE epitope (P) at the N-terminus (MB1N, MB2N, MB4N) or at the N- and C-terminus (MB1N1C) along with a C-terminal 6xHistidine tag (H). Glycine spacers between the IgE-epitopes are shown as black bars. (*B*) Tertiary structure of MB1N obtained by mathematical modelling. The IgE epitope at the N-terminus is shown in red. (*C*) Surface plasmon resonance determination of the MB1N-IgE interaction by immobilization of IgE and injection of increasing concentrations of MB1N. Signal intensities (response units, RU, y-axis) were measured over time. Recorded data (black) and calculated curves (red), which represent fittings of the data to a 1:1 binding model, were superimposed. Association rate constant (ka), dissociation rate constant (kd) and dissociation constant (KD) are indicated. (D-G) Modelling of tertiary structures of MB1N1C (D, open conformation; E, closed conformation), MB2N (F), and MB4N (G). Coloring of epitopes corresponds to the scheme in (A).

Using the protein structure prediction algorithms RosettaFold and AlphaFold 2, we generated representations of the 3D structures of the myoglobin backbone plus its derivatives. 3D models of MB1N, MB2N, MB1N1C, and MB4N are shown in Figs. 1 C-G. The structure of the monovalent form, MB1N (Fig. 1C) showed that myoglobin is organized into 8 α-helical regions (gray), with the single Phl p 1 epitope in red added at the N-terminus forming a fully accessible alpha helical structure. For MB1N1C, modelling predicted 2 spatial configurations. The first is an open structure (Fig. 1D), where the two IgE epitopes are both surface exposed and in greater distance from each other than in MB2N, and a closed form (Fig. 1E), where the C terminal epitope is hidden in a pocket formed by the myoglobin helices. As both structures were minimized to very similar energy values, MB1N1C may exist in solution as a mixture of mono- and bivalent molecules. For MB2N and MB4N, either two or four Phl p 1 epitopes extend from the N-terminus in repeating alpha helical structures. Both MB2N and MB1N1C contain two grafted epitopes. In tertiary structure, the Euclidean distance between the two epitopes are similar with 20.88 Å for MB2N, 18.47 Å for MB1N1C (closed), and 24.14 Å for MB1N1C (open).

### Higher valency antigen elicits stronger secretory responses than bivalent antigen

Consistent with previous studies^20^, we found that the higher valency MB4N was more efficient at eliciting a degranulation response than either bivalent antigen. Figure 2A shows the degranulation response in the well-known rat basophilic leukemia (RBL-2H3) cell model. Cells were primed with a saturating dose of anti-Phl p 1 IgE (2 μg/mL mIgE for 2 h) and challenged with MB1N, MB2N, MB1N1C, or MB4N over a range of antigen doses (0.01-10 nM). Notably, MB4N induces degranulation at low concentration of only 0.1 nM, while MB2N and MB1N1C do not elicit a response until the intermediate dose of 1 nM antigen. At the high dose of 10 nM, all antigens show a robust response.

**FIGURE 2.**
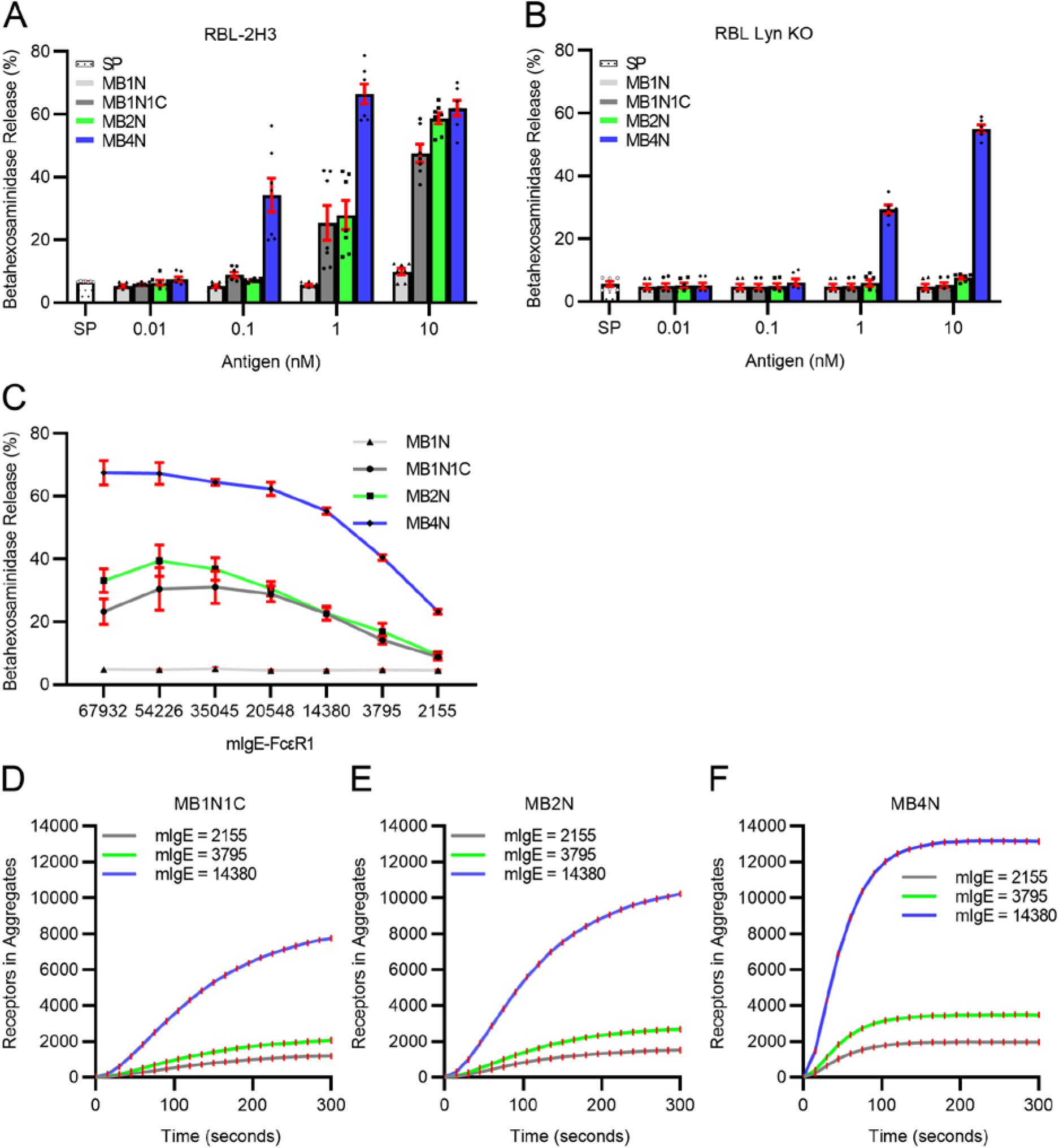
Differential receptor aggregation and degranulation response dependent on antigen concentration and valency, FcεRI occupancy by IgE antibodies, and availability of Lyn kinase. (A) RBL-2H3 cells and (B) RBL-Lyn^KO^ cells loaded with the peptide-specific monoclonal IgE antibody were incubated with allergen constructs MB1N, MB2N, MB1N1C, or MB4N at the indicated concentrations (x-axis). Beta-hexosaminidase release is displayed as percentage of total release (y-axis). Mean values ± SEM from 6 wells. SP, spontaneous release without antigen. (C) RBL-2H3 cells were primed with different concentrations of mIgE. Numbers of occupied FcεRI as determined by FACS measurement are given at the x-axis. Degranulation in response to 0.1 nM MB1N, MB1N1C, MB2N, and MB4N is shown at the y-axis as % ß-hexosaminidase release. Plots show mean ± SEM for four experiments. (D-F) Rule-based model simulation of FcεRI aggregation over time showing the number of receptors in aggregates induced by (D) MB4N, (E) MB2N, and (F) MB1N1C. Values are mean ± SEM.

We sought further insight into the influence of antigen valency upon mast cell degranulation by comparing results in RBL cells lacking Lyn tyrosine kinase through CRISPR/Cas 9 gene editing (RBL-Lyn^KO^; Kanagy et al^23^), primed with saturating doses of anti-Phl p1 mIgE. Results in Figure 2B illustrate the dramatic differences between the bivalent and tetravalent antigens. MB4N (blue bars, Fig. 2B) can induce degranulation without Lyn, albeit needing higher dose than in parental cells (Fig. 2A). In contrast, MB2N (green bars) and MB1N1C (dark gray bars) cannot support degranulation in RBL-Lyn^KO^ cells at any dose. These results confirm that higher valency antigens can override the need for Lyn-mediated phosphorylation of receptor ITAMs, until recently considered to be the earliest essential step in the canonical FcεRI cascade^23^.

Since allergic patients vary widely in antigen-specific IgE isotypes and their relative abundance, we next investigated the impact of the number of receptors occupied with mIgE on the secretory response of RBL-2H3 cells to the structurally defined ligands. To this end, a flow cytometry-based assay was used to quantify the occupancy of FcεRI on the cell surface with monoclonal IgE antibodies. RBL-2H3 cells were exposed to fluorescently labeled mIgE for 2 h at concentrations ranging between 10 and 600 ng IgE/mL. These priming conditions translated to an occupancy of ∼2000 up to ∼68,000 IgE-bound receptors on the cell surface (Supplementary table S1; X axis, Figure 2C). Figure 2C shows results from a degranulation assay after 2 h priming with these IgE concentrations and activating with a 1 nM concentration of the different ligands or myoglobin as a negative control. MB4N induced the highest ß-hexosaminidase release at all IgE concentrations, achieving maximal response at ∼35,000 occupied FcεRI (63% secretion). Remarkably, an IgE receptor occupancy of ∼2000 was sufficient for a 25% secretory response to MB4N activation. Both MB2N and MB1N1C were less effective in inducing degranulation at all IgE priming conditions, compared to MB4N. Maximal release was achieved for MB2N (48%) and MB1N1C (42%) when 35,000 to 54,000 receptors were occupied and dropped to 11% for both ligands with only 2000 occupied receptors. Differences in allergenic activity between MB2N and MB1N1C, which were observed in previous studies, only occurred at high IgE:receptor numbers^20^. Neither the scaffold protein myoglobin nor the monovalent version (MB1N) induced degranulation at any IgE concentration (results not shown).

A rule-based model simulation of aggregate formation on the cell surface was performed for MB1N1C (Fig. 2D), MB2N (Fig. 2E), and MB4N (Fig. 2F). After the first 5 minutes since exposure, the number of receptors in aggregates stabilizes for MB4N, but is still increasing for MB2N and MB1N1C. The final number of receptors in aggregates is directly proportional to the number of available receptors in each simulation. For each of the mIgE numbers, MB4N has consistently a larger number of receptors in aggregates due to its higher valency (Figs. 2D-F).

#### Calcium signaling downstream of FcεRI crosslinking is highly sensitive to antigen valency

A rise in intracellular calcium levels is essential to antigen-stimulated mast cell secretion ^35,36^. We next used single cell imaging to measure responses to the bivalent and tetravalent antigen. Calcium flux was monitored in parental RBL-2H3 cells preloaded with the esterified, ratiometric calcium dye, Fura-2^25,37^. Heat maps in Figures 3A-C report the range of calcium responses in dozens of individual cells, where each row indicates the change in Fura-2 ratio over time for a single cell. Consistent with robust degranulation, crosslinking by MB4N typically induced a rapid and sustained intracellular calcium release (Figure 3C). A minor fraction of cells (∼15%, blue bars in Figure 3D) triggered with the tetravalent antigen were unresponsive at the highest dose of 1 nM; at the lower dose of 0.1 nM, approximately 75% of cells responded with a measurable rise in calcium. In contrast, large fractions of cells treated with either of the bivalent antigens failed to trigger responses (gray and green bars in Figure 3D). In those cells where calcium responses were recorded by MB2N (Fig. 3B) or MBN1C1 (Fig. 3A), the response was delayed and lower in magnitude than for the higher valency antigen. Significantly shorter lag times (Figure 3E,G) and larger overall response (rise height in Fig. 3F,H) were also observed for the tetravalent antigen, compared to both bivalent antigens.

**FIGURE 3.**
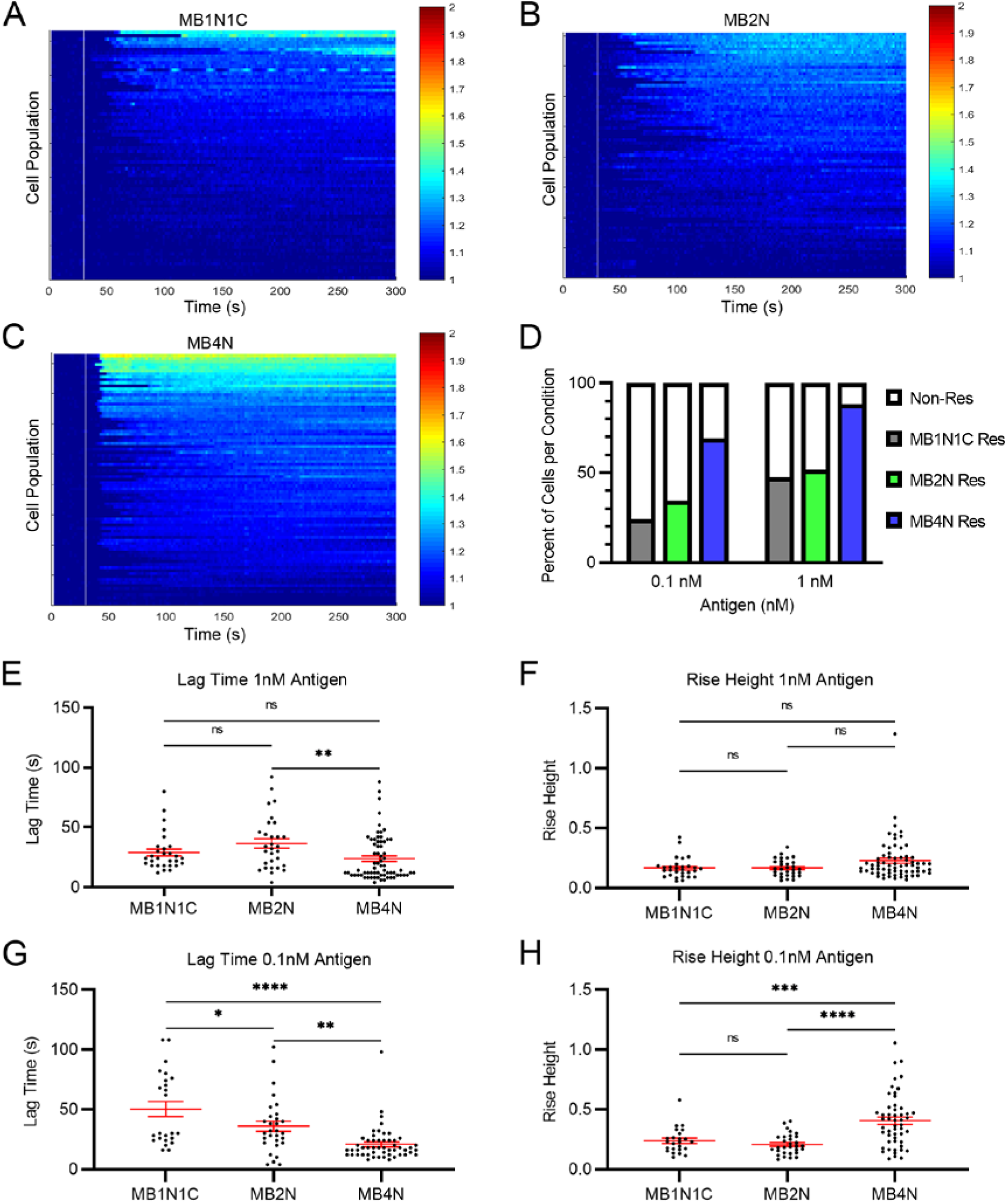
Calcium release in mIgE-primed RBL-2H3 cells after activation with MB1N1C, MB2N, or MB4N. (A-C) Heatmaps show relative changes in intracellular calcium concentration after addition of 1 nM (A) MB1N1C, (B) MB2N, or (C) MB4N at 30 sec (vertical white line). Each row represents the ratio of Fura-2 emission using 340nm/380nm excitation over time for a single cell. (D) Percentage of cells responding to MB1N1C, MB2N, and MB4N at the indicated concentrations. Number of cells analyzed in each condition ranged from 58 to 99. (E, G) Time between addition of (E) 1 nM antigen or (G) 0.1 nM antigen and the response for each cell. (F, H) Relative increase in Fura-2 ratio per cell after addition of (F) 1 nM antigen or (H) 0.1 nM antigen. Mean values and SEM are indicated. * P < 0.05, ** P < 0.01, *** P < 0.001, **** P < 0.0001, ns, not significant. Data in E-H reports only on cells that responded to antigen; each dot indicates a single cell measured.

### Valency dictates aggregate size and complexity

Previous work with the Phl p 1-derived antigen variants used transmission electron microscopy (TEM) to illustrate the different types of aggregates that can be formed by crosslinking of IgE in solution^20^. We next used a computational model (Fig. 4) to estimate the geometry of FcεRI-IgE aggregates, when constrained by orientation on the cell surface after exposure to bivalent and tetravalent Phl p1-derived antigens. Example aggregates are shown for MB1N1C, MB2N and MB4N in Figures 4A-C, respectively (see also Supplemental Movies 1-3). From our Monte Carlo simulations, we can measure aggregate size based on 2 metrics: Number of receptors per aggregate (Fig. 4D), and aggregate spread, measured in angstroms from the center of mass of the aggregate (Fig. 4E). Our results show that MB4N and MB2N have, on average, the same number of receptors per aggregate, with MB4N presenting more outliers, and are both larger than aggregates formed with crosslinking via MB1N1C (Fig. 4D). In terms of aggregate spread, MB4N aggregates are significantly larger than MB2N and MB1N1C (Fig. 4E). These results are consistent with antigen valency increasing aggregate size.

**Figure 4.**
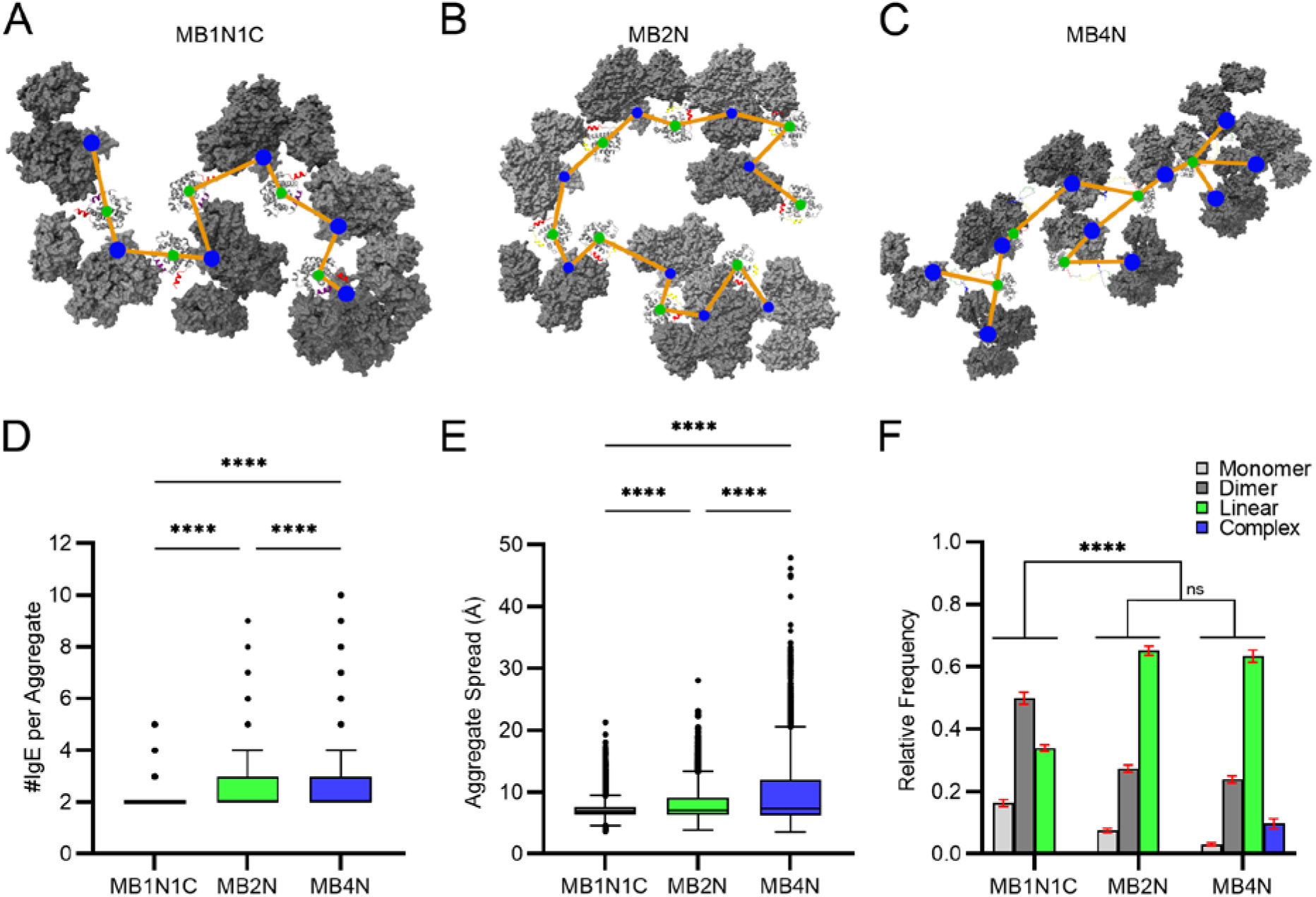
Results from Monte Carlo simulation of IgE-Antigen aggregation. (A-B) Depiction of examples of linear aggregates formed with MB1N1C and MB2N, both valency 2 allergens. See also Supplemental Videos 1-3. (C) A complex aggregate formed by crosslinking of IgE via MB4N. Complex aggregates can only be formed when valency is greater than 2. (D) Number of IgE molecules per aggregate. MB1N1C contains fewer IgE molecules on average than MB2N and MB4N. (E) Aggregate spread in angstroms for each allergen model, with MB4N forming larger aggregates in size than MB1N1C and MB2N. (F) Formation of aggregates per model. All models are able to form linear aggregates, with MB2N and MB4N forming a similar ratio of linear chains that is larger than the ratio of linear aggregates formed by MB1N1C. MB4N is the only model capable of forming complex aggregates, observed in nearly 10% of all aggregates formed during the course of the simulation. Data from D, E represents results from >15,000 simulations and F is from 100 simulations. **** p<0.0001, ns is not significant; two-way ANOVA Tukeys multiple comparisons test.

Simulation results show that the bivalent MB2N antigen bound to bivalent IgE-FcεRI primarily generates dimers and short linear chains, with a higher ratio of the latter (Fig. 4F). Bivalent MB1N1C also generates dimers and linear chains, but with a higher proportion of dimers due to the occlusion of the C-terminus in half the molecules (Fig. 4F). The example shown in Figure 4A (see also Supplemental Video 1) shows a linear chain of MB1N1C with 5 antigens and 6 receptors. Figure 4B illustrates a chain of 8 receptors engaged with 8 bivalent MB2N ligands (see also Supplemental Video 2). In contrast to results obtained in solution^20^, we did not observe the formation of any cyclic structures, suggesting that the membrane orientation of the complexes places constraints that prevent cyclic aggregates. By comparison, the tetravalent MB4N generates highly branched and complex aggregates, in addition to dimers and linear chains (Fig. 4F). In Figure 4C, we show a simulated complex aggregate of 10 receptors bound to 5 MB4N antigens (see also Supplemental Video 3).

Monte Carlo-based simulation including tertiary structures of MB2N and MB1N1C provides insight into steric hindrance that limits epitope accessibility. This is best seen by comparing conditional binding probabilities, P(N|C) and P(C|N), of epitopes N (N-terminus) and C (C-terminus), on a single antigen. For example, MB2N shows that binding either epitope similarly impacts the other as seen in the conditional probabilities, P(N|C) = 0.60 and P(C|N) = 0.56. On the other hand, MB1N1C (closed) demonstrates that the N-terminus epitope more strongly influences binding on the C-terminus epitope (probability P(C|N) = 0.27), while the C-terminus epitope has less influence on binding the N-terminus epitope (probability P(N|C) = 0.71). The same is true for MB1N1C (open) with conditional binding probabilities P(C|N) = 0.48 and P(N|C) = 0.77.

Model predictions were validated experimentally using two high resolution microscopy approaches. Figure 5A shows an example TEM image of membrane sheets^22^ ripped from parental RBL-2H3 cells primed with anti-Phl p1 mIgE but not exposed to antigen; IgE receptors are marked by sequential labeling with anti-FcεRI-β antibodies and anti-mouse IgG conjugated to 6 nm colloidal gold. As expected based upon prior work^22^, resting FcεRI are dispersed across the cell surface as singlets and very small clusters. This slightly non-random distribution is confirmed by rightward shift of the Gaussian fit (green line) for data acquired and analyzed using the Hopkins test (Fig. 5E, top)^38^. Prior work^39^ supports a model where this non-random distribution observed in fixed cells is due to transient residency of mobile IgE receptors in microdomains. Results in Figure 5B-D show the impact of antigen stimulation on receptor distribution, with a visual confirmation of an increase in small chains and clusters after treatment with the bivalent antigens, MB1NC and MB2N; the Hopkins test shows an atypical biphasic shift in the right-shifted Gaussian fit that potentially reflects the mix of short chains and small clusters. Visual inspection of FcεRI distribution after treatment with the tetravalent antigen, MB4N, shows larger clusters and a marked rightward shift in the Hopkins test. The representative image in Figure 5D has two large clusters in the lower right.

**FIGURE 5.**
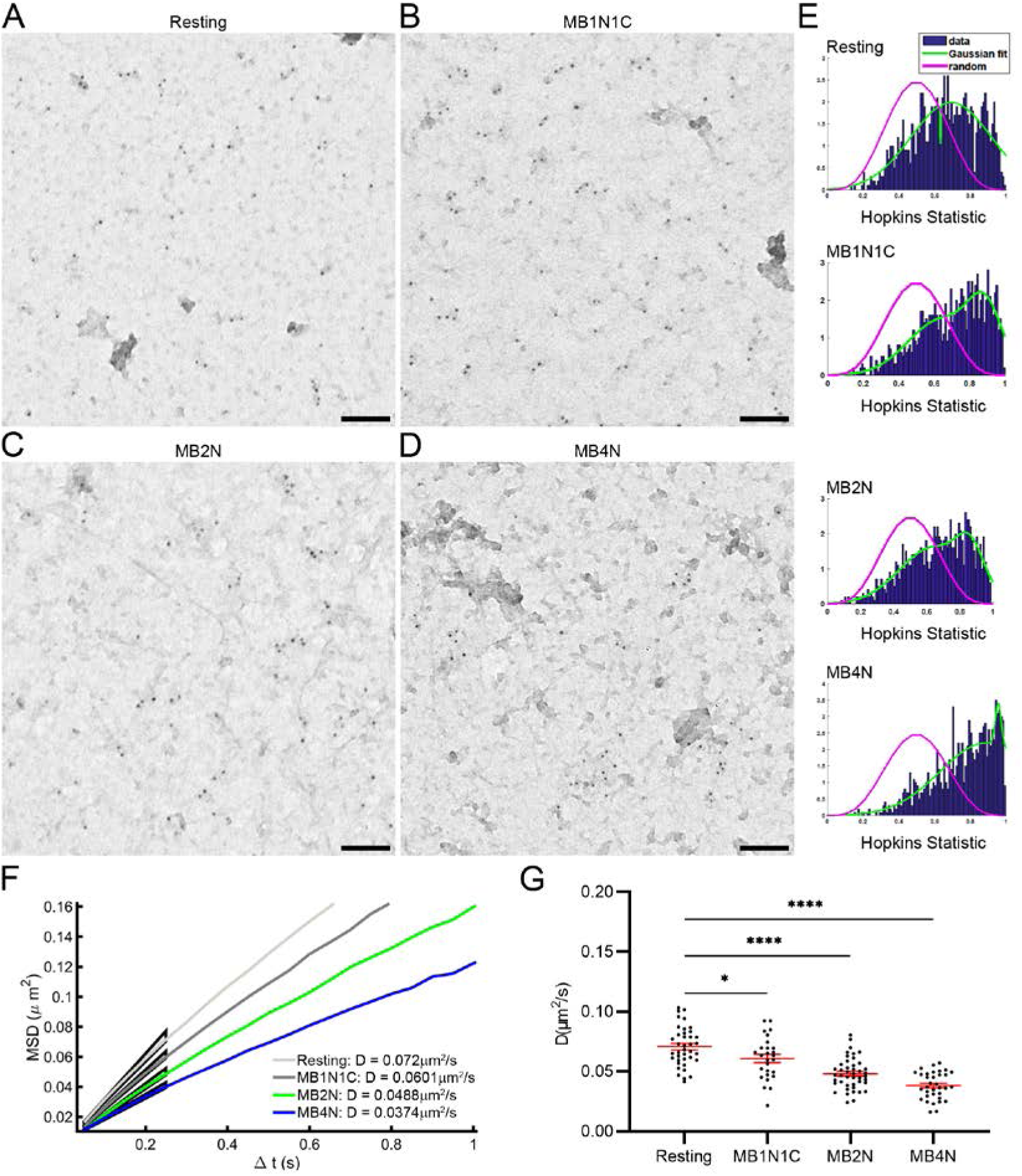
Antigen-dependent FcεRI aggregation and diffusional dynamics on the surface of RBL-2H3 cells. (A-D) TEM images of membrane sheets prepared from cells primed with peptide-specific mIgE, treated with buffer (A, Resting), or stimulated with (B) MB1N1C, (C) MB2N, or (D) MB4N and immunogold labeled for FcεRIβ. Scale bar, 100 μm. (E) Evaluation of conditions in A-D by Hopkins statistics. (F) Single particle tracking of HA-QD labeled FcεRI was used to quantify receptor mobility on RBL cells when crosslinked with different antigens. Ensemble MSD calculated across all cells of single condition. Reduction in slope of the MSD indicates reduced mobility. Fitting the MSD returns the Diffusion coefficient (D), reported for each condition in the legend. (G) Corresponding distribution of ensemble diffusion coefficients, D, calculated for each individual cell (black circle), showing the spread in D; * p < 0.05,** p < 0.01,*** p < 0.001. Red bars indicated mean ± SEM. n > 27 cells analyzed per condition.

Single particle tracking (SPT) is a sophisticated method to track changes in motion for individual IgE receptors after crosslinking with multivalent antigens^39^. In a significant technical advance^27^, SPT data can now be acquired in RBL cells stably expressing recombinant FcεRI γ-subunits with an extracellular HA tag that are labelled with SPT without the need to modify the anti-Phl p1 mIgE used for priming. Figure 5 reports SPT data acquired in resting cells versus cells stimulated with bivalent or tetravalent antigens, shown as ensemble mean squared displacement (MSD) across all cells of a single condition (Fig. 5F) or as the average D value per cell (Fig. 5G). The light gray line in Fig. 5F reports the typical rapid diffusion behavior of resting IgE receptors, with a diffusion coefficient (D) value of 0.072 µm^2^, as calculated from the MSD. Receptors crosslinked with MBN1C1, which is likely fluxing in solution between closed and open forms (Fig. 1D,E), exhibit a modest slow down with an MSD value of 0.0601 µm^2^. The stable bivalent MB2N antigen slows IgE receptors more notably, with D = 0.0488 µm^2^. The tetravalent MB4N antigen markedly slows receptors upon crosslinking, with an MSD value of 0.0374 µm^2^, consistent with the larger aggregates predicted by modeling and observed by TEM on membrane sheets (Figs. 5D, F).

## Discussion

In this study we used relevant molecular model antigens to study FcεRI aggregation on the cell surface of mast cells and subsequent intracellular events. The engineered molecules represent relatively small protein antigens, as observed for most allergens recognized by IgE antibodies in allergic patients^1^. They are based on an important grass pollen allergen-derived peptide sequence, which binds to the corresponding monoclonal IgE with high affinity, comparable to allergen-specific IgE cloned from allergic patients^40^. The model antigens differ in epitope number and spacing, factors which were previously shown to play a role in *in vitro* and *in vivo* models of mast cell and basophil activation^20,21^. The present work combines state-of-the-art imaging approaches with computer simulations to correlate the states of FcεRI aggregation on the cell surface with signaling efficiency. Novel insight is also provided by comparing degranulation responses after varying FcεRI occupancy with epitope-specific IgE prior to crosslinking with model antigens of different valencies. Overall, results confirm that each of these variables (receptor occupancy, antigen valency and antigen concentration) markedly impacts mast cell responses.

Monte Carlo-based simulations of allergen-induced receptor crosslinking on the cell surface were uniquely applied here to predict the structural complexity and size of FcεRI aggregates. MB2N and MB1N1C represent a class of allergens with a valency of two, either through epitope repeats or homodimerization^41^. Wofsy and Goldstein^42^ proposed that bivalent antigen antibody complexes can only form chains or closed rings. These configurations were both observed by TEM for MB2N- IgE complexes formed in solution^20^. However, our modeling results would suggest that steric constraints imposed by tethering to the plasma membrane led to a predominance of linear chains for both bivalent antigens used here. Only the tetravalent MB4N seems to be able to induce larger, branched aggregates.

Computer simulations support the observed aggregation on membrane sheets analyzed by TEM, as well as receptor mobility changes observed in SPT experiments where the diffusion coefficients are ordered from slowest to fastest: 0.037 μm^2^/s, 0.049 μm^2^/s, and 0.060 μm^2^/s, for MB4N, MB2N, and MB1N1C, respectively (Fig. 5F,G). In simulation, MB4N produces aggregates of similar receptor number as MB2N but the aggregates it produces have larger spread (median=25 Å) than either of the bivalent antigens (median=20 Å, Figure 4E). Crosslinking efficiency of MB4N is likely influenced by its higher valency and the steric flexibility of the four epitopes at the N-terminus of the protein (Figure 1G).

Two different tertiary structures were predicted by models for MB1N1C, which differ in accessibility of the IgE epitope and may be interchangeable in solution. While these predictions would require validation by crystallization or NMR, the potential for differential availability of IgE epitopes introduces interesting concepts that may be relevant to natural allergens that modulate between different states. Rouvinen and colleagues have shown that lipocalin allergens can form transient dimers and propose that this may apply to other classes of allergens^41,43^.

For several decades, the consensus view has been that FcεRI signalling is both initiated and downregulated by Lyn-mediated phosphorylation of FcεRI β and γ ITAM motifs^44,45^. Recent development of new tools, such as RBL cells lacking Lyn through CRISPR/Cas 9 gene editing, have led to the emerging concept that differences in aggregate formation translate to differential signaling at very early steps in FcεRI signaling. Using RBL-Lyn^KO^ cells, Kanagy et al showed that high valency ligands, such DNP_25_-BSA targeting receptors bound to DNP-specific IgE, can override the need for Lyn in initiating FcεRI signaling^23^. The current study expands this observation by showing that the tetravalent antigen, MB4N, also overcomes Lyn deficiency. In contrast, activation by the bivalent antigens remain completely reliant on Lyn. The role of aggregate geometry in effector cell activation clearly merits further study.

We observed a reduced degranulation of RBL-2H3 cells when fewer FcεRI receptors were occupied by allergen-specific IgE but find that low receptor occupancy can be overcome by the higher valency antigens that lead to bigger aggregates on the cell surface. This is in line with *in vitro* studies in primary basophils and mast cells ^21,46,47^.

These findings translate to clinical observations at several different levels. Early studies already revealed that circulating IgE antibodies to ragweed correlated with those tightly bound to FcεRI on blood basophils. Moreover, a significant relationship between IgE levels specific for ragweed allergens, which increased after allergen exposure, and disease severity was observed^48^. Using defined monoclonal IgE antibodies and purified allergenic molecules, the correlation of allergen-specific IgE levels and effector cell degranulation was confirmed *in vitro* in basophils and mast cells ^21,46,47^. Furthermore, clinical data suggest that high levels of IgE to certain allergens, like the major grass pollen allergens, Phl p 1, Phl p 2, Phl p 5, and Phl p 6, and the major peanut allergen, Ara h 2, associate with severe allergic symptoms^49–51^. Importantly, a positive correlation between IgE epitope diversity (i.e., number of epitopes recognized) and clinical sensitivity was also found in peanut and milk allergic children^52,53^. Structural features of allergens, including those where substantial flexibility occurs around epitopes, may also be a fruitful new focus area for understanding the potency of the molecule in activating effector cells, as has recently been described for the major grass pollen allergen, Phl p 5^54^. Therefore, the described model represents a realistic depiction of the processes in allergic patients allowing the detailed dissection of factors involved in immediate allergic reactions.

## Acknowledgements

This work was funded by grant NIH R35GM126934 (DSL), the New Mexico Spatiotemporal Modeling Center NIH P50GM085273, the Danube Allergy Research Cluster program funded by the Country of Lower Austria and the National Science Foundation IIS-1553266 (LT). JCD was supported in part by a Department of Education GAANN award. We thank Dr. Cedric Cleyrat and Eunice Choi for generation of the knockout cell lines. We gratefully acknowledge use of the University of New Mexico Comprehensive Cancer Center fluorescence microscopy and flow cytometry facilities, as well as NIH-NCI support via P30CA118100 for these cores. We would like to thank the UNM Center for Advanced Research Computing, supported in part by the National Science Foundation, for providing the high-performance computing resources used in this work.

## Supplementary Video Legends

**Video S1.** Model of a linear aggregate formed by MB1N1C. Linear aggregates consist of alternating antibody-ligand links with no branching. Here, ligand in a linear aggregate is bound to at most two antibodies. Antigen: Gray PDB model of MB1N1C with epitopes identified according to the scheme in Figure 1A. Antibodies: Gray 3D isosurface model of IgE bound to FcεR1α subunit.

**Video S2.** Model of a linear aggregate formed by MB2N. Linear aggregates consist of alternating antibody-ligand links with no branching. Here, ligand in a linear aggregate is bound to at most two antibodies. Antigen: Gray PDB model of MB2N with epitopes identified according the scheme in Figure 1A. Antibodies: Gray 3D isosurface model of IgE bound to FcεR1α subunit.

**Video S3**. Model of a complex aggregate formed by MB4N. Complex aggregates consist of alternating antibody-ligand links with branching. Consequently, there exists at least one ligand in a complex aggregate that is bound to at least three antibodies. Antigen: Gray PDB model of MB4N with epitopes identified according to the scheme in Figure 1A. Antibodies: Gray 3D isosurface model of IgE bound to FcεR1α subunit.

---

**Supplementary Table S1.** Quantification of mIgE-bound FcεRI on RBL-2H3 cells.

| mIgE<sup>Alexa647</sup> concentration for priming of RBL cells (ng/ml) | No. of calculated FcεRI bound to mIgE<sup>Alexa647</sup> on cell surface |
| --- | --- |
| 600 | 67932 |
| 300 | 54226 |
| 150 | 35045 |
| 75 | 20548 |
| 38 | 14380 |
| 19 | 3795 |
| 10 | 2155 |

